# Novel carbon nanoparticles derived from *Bougainvillea* modulate vegetative growth in *Arabidopsis*

**DOI:** 10.1101/2023.06.19.545555

**Authors:** Raviraj B Barot, Nilesh D Gawande, Satya Omprabha, Charli Kaushal, Jhuma Saha, Dhiraj Bhatia, Subramanian Sankaranarayanan

## Abstract

We present a green synthesis method of producing blue fluorescence emitting carbon nanoparticles (CNPs) through a simple and cost-effective single-step hydrothermal reaction. The synthesis utilized bract extracts and pollen grains from three *Bougainvillea* species: *B. spectabilis, B. alba*, and *B. buttiana*. The CNPs exhibited photoluminescence, with the highest emission observed in the ultraviolet region. Atomic force microscopy analysis revealed that the size of synthesized CNPs ranged from 23 nm to 83 nm. Fourier transform infrared analysis provided a comprehensive understanding of the CNP’s surface functional groups, with carbon being the predominant group. X-ray diffraction analysis confirmed the amorphous nature of the synthesized CNPs. Zeta potential measurements indicated that the particles carried a negative charge, suggesting their colloidal stability. In experiments conducted with *Arabidopsis thaliana* seedlings, CNPs derived from *B. alba* pollen grains were found to promote leaf area expansion while simultaneously inhibiting primary root growth. Conversely, other CNPs demonstrated detrimental effects on vegetative growth. These findings underscore the potential application of these novel CNPs in agriculture.

## Introduction

Carbon, the most abundant mineral on the Earth, serves as the fundamental building block for essential macromolecules that support life, such as carbohydrates, proteins, DNA, and other vital components. Pure carbon exists in the form of several allotropes, with diamonds and graphite being the most commonly found naturally occurring forms. Conversely, amorphous carbon allotropes, including coal, charcoal, and lampblack, also exist (Kroto et al. 1985; Sharma 2010). Nanotechnology has revolutionized the synthesis of a broad range of materials at the nanoscale level. Nanoparticles (NPs), which encompass particulate materials with a diameter smaller than 100 nm, form a significant category within this realm (Khan et al. 2019). Among the diverse types of nanoparticles, carbon nanoparticles (CNPs) were first discovered in the 1980s (Kroto et al. 1985; Sharma 2010). CNPs encompass a wide array of carbon-based structures, including amorphous CNPs (such as ultrafine carbon particles, carbon nanoparticles, and carbon dots), sp2 carbon nanomaterials (such as fullerene, carbon nanotubes, carbon nanohorns, graphene, and graphene quantum dots), and nanodiamonds (Gaur et al. 2021). The CNPs have distinct chemical and optical characteristics and can be synthesized using a variety of processes, including electrochemical and physical procedures, chemical vapor deposition (CVD), high-radiation-band preparation, thermal deposition, laser ablation, the oxidation of C materials, and the chemical disintegration of fiber and carbohydrates (Campuzano et al. 2019; Zhao et al. 2019; Sharma and Das 2019; Guo and Zhao 2020).

CNPs, with a size of approximately 100 nm, have emerged as a versatile tool for various applications, including medication administration, gene therapy, medical imaging, cancer treatment, and many more applications (Ray and Jana 2017). In the realm of biomedical applications, CNPs hold immense promise for tasks such as biological labelling, bioimaging, and other optoelectronic/electronic device applications. However, the utilization of green biosynthesis techniques may offer significant advantages when synthesizing CNPs from natural sources (Ray and Jana 2017; Guerrero et al. 2017). The synthesis of CNPs using biomass as a carbon source has the potential to be both economically and environmentally advantageous, in contrast to chemical carbon sources (Sharma et al. 2019).

CNPs show promising applications in plant growth regulation. Various studies have explored the effect of nanoparticles derived from different sources on plant growth and abiotic stress management (García-Sánchez et al. 2015; Kumar et al. 2018; El-Shetehy et al. 2021; Singh et al. 2021). For instance, exposure to copper nanoparticles has been found to adversely affect the plant agronomic and physiological characteristics of cilantro plants, resulting in significant reductions in germination, shoot elongation, chlorophyll, and nutrient accumulation (Zuverza-Mena et al. 2015). Wheat plants exposed to a concentration of 15.6□μM copper nanoparticles displayed a remarkable 60% decrease in relative root growth, while lateral root growth was stimulated at similar conditions (Zhang et al. 2018). Similarly, exposure to the 10–150 µg/ml of MgO NPs had a detrimental impact on the morphological and biological characteristics of black gram in a concentration-dependent manner, resulting in a decrease in shoot-root length and the shoot/root ratio (Sharma et al. 2021). Exposure to nanoparticles can also induce transcriptomic changes in the model plant *A. thaliana* (Landa et al. 2012).

In contrast, numerous studies have demonstrated the beneficial effect of nanoparticles. For example, nanoparticles derived from AuNPs have been shown to possess larvicidal and nematicidal properties in crop cultivation, effectively controlling pests without harming crop growth and development (Sundararajan and Ranjitha Kumari 2017). The AuNP-SCTAs derived nanoparticles had a favourable effect on the growth of Arabidopsis seedlings, safeguarding them against oxidative stress by eliciting immunological responses (Ferrari et al. 2021). Transcriptomics and proteomics studies supported the hypothesis that the trade-off between growth and immunological/stress responses shifts towards favouring growth following AuNP exposure. These studies revealed the downregulation of oxidative stress, and immunological responses, along with the overexpression of growth-promoting genes (Ferrari et al. 2021). Furthermore, the combination of TiO2 and SiO2 nanoparticles increased the activity of nitrate reductase in soybean, facilitating faster germination and growth through efficient resource acquisition, including water and fertilizers (Lu et al. 2015). Similar growth-promoting effects through resource utilization were also observed in mung bean (Li et al. 2016). Collectively, these studies provide comprehensive evidence of the effect of nanoparticles on plants at various levels, such as the transcriptome, proteome, and phenotypic levels, indicating both the stimulatory and inhibitory effects of nanoparticles on plant growth and development.

The *Bougainvillea* genus, belonging to the Nyctaginaceae (4 O’clock) family, encompasses approximately 18 species. Among these species, only four species (*B. buttiana, B. glabra, B. spectabilis*, and *B. peruviana*) have been commercially utilized (Abarca-Vargas and Petricevich 2018). Extensive research has been conducted on the chemical composition of various parts of *Bougainvillea*. While primarily recognized as an ornamental plant, *Bougainvillea* holds a significant medicinal value. Numerous studies have demonstrated its anticancer, antimicrobial, antioxidant, anti-inflammatory, and antihyperglycemic properties (Chauhan et al. 2016; Ogunwande et al. 2019; Ahmar Rauf et al. 2019). Certain *Bougainvillea* species, like *B. spectabilis*, also provide benefits for individuals with diabetes and are also employed in the treatment of coughs and sore throats, hepatitis, joint pain, fever, and ulcers, and as an effective body detoxifier (Guerrero et al. 2017; Abarca-Vargas and Petricevich 2018). Moreover, laboratory testing has revealed that an extract from the leaves of *B. spectabilis* exhibits reduced tospovirus, the causative agent of spotted wilt on *Capsicum annum* in the groundwater (Balasaraswathi et al. 1998). These findings underscore the potential of different *Bougainvillea* species in the treatment of various diseases, warranting further exploration of their effects in other domains.

In this study, we synthesized novel CNPs through a single-step hydrothermal reaction, utilizing bract extract and pollen grains of *Bougainvillaea* species. We further characterized these CNPs using various approaches. The size of these CNPs ranged from 23 nm to 83 nm. The synthesized CNPs exhibited amorphous nature and carried a negative charge. When exposed to UV light, these CNPs emitted a bluish-purple fluorescence. FTIR spectroscopy analysis revealed the presence of diverse functional groups on the surface of the CNPs. Interestingly, the synthesized bract extract CNPs displayed similar functional groups, indicating variations in characteristics and functions between the two types of CNPs derived from different sources. We also carefully examined the effect of CNPs on *A. thaliana* seedling growth. Our study suggests that CNPs synthesized from different *Bougainvillea* species influence the vegetative growth of *Arabidopsis* plants in contrasting ways.

## Materials and methods

### Synthesis of Carbon nanoparticles

Bracts and pollen grains from three different species of *Bougainvillea* (*B. spectabilis, B. alba*, and *B. buttiana*) were collected from the Indian Institute of Technology Gandhinagar (IIT-GN) campus, Gujarat, India. The bract extract for the *Bougainvillea* species was prepared using the protocol described by Sharma *et al*. (2019). The bract extract was mixed with 300 mg L-cysteine (Himedia) and 300 µL of ethylenediamine (Sigma-Aldrich) for 10 minutes using a magnetic stirrer, and this mixture was heated in a Teflon-lined hydrothermal autoclave at 200°C for 5 hours. The reactor was allowed to cool naturally, and the dark brown solution was collected. The solution was centrifuged for 10 minutes at 10,000 rpm and 25°C, and the supernatant was collected and filtered using a 0.22 µm membrane filter and stored at room temperature for further use (**Supplementary Fig. S1A**)

A small quantity of 0.1 gram of pollen grains was carefully collected from three *Bougainvillea* species, namely *B. spectabilis, B. alba*, and *B* .*buttiana* species, by dissecting the stamen. The collected pollen grains were then suspended in 30 mL of ultrapure water, followed by sonication for 15 minutes to ensure proper dispersion. Subsequently, this mixture was placed in a Teflon-lined hydrothermal autoclave and heated at 200 □ for 20 hours. After completion, the reactor was allowed to cool down to room temperature, and the resulting solution was filtered using a 0.22 µm membrane filter and stored at room temperature (**Supplementary Fig. S1B**). To distinguish between carbon nanoparticles synthesized from bract extract and those derived from pollen, the former were denoted as Bs-PET, Ba-PET, and Bb-PET, representing *B. spectabilis, B. alba*, and *B. buttiana*, respectively. On the other hand, the latter were labelled as Bs-POL, Ba-POL, and Bb-POL for the corresponding *Bougainvillea* species.

### Plant growth conditions and vegetative growth assay

Seeds for the *Arabidopsis* wild-type genotype *Col-0* were sterilized for 5 min with 70% ethanol, followed by 4-5 times wash with sterilized distilled water. Seeds were germinated on the sterile half Murashige Skoog (MS) agar media plates containing 0.22% MS salt, 0.05% MES monohydrate, 2% sucrose, and 0.8% plant agar, and the pH of the media was adjusted to 5.7 using 1M KOH. Plates were incubated in the dark at 4°C for two days and transferred to the plant growth chamber at 22°C with a light intensity of 130 µmol/m2/s under long-day conditions (16 hr light / 8hr dark). Three-day post-germination seedlings were transferred to the control MS plates (without CNPs supplementation) and MS plates supplemented with three different concentrations of CNPs (2.5mg/ml, 5mg/ml, and 10mg/ml) for Bs-PET, Ba-PET, Bb-PET, Bs-POL, Ba-POL, and Bb-POL. The effect of the nanoparticles on the root growth and leaf area was quantified on day seven after the transfer of the seedling using Image J.

### Characterization of the *Bougainvillea* pollen morphology and CNPs

The morphology of pollen grains from three *Bougainvillea* species was determined using a field emission scanning microscope (FE-SEM, JEOL® JSM-7600F). CNPs were extensively characterized using different methods. The structural morphology of CNPs was determined using an atomic force microscope (AFM, Bruker® Nano wizard Sense AFM). The functional group in CNPs was identified using a Fourier-transform infrared spectrometer (FTIR, Perkin Elmer® Spectrum Two FTIR Spectrometer). The nature of the CNPs was determined using multipurpose X-ray diffraction (M-XRD, Rigaku® SmartLab 9KW). The optical properties of the CNPs were analyzed using a UV-VIS spectrophotometer (Analytika Jena SPECORD® 210 PLUS) and a spectrofluorometer (Jasco® Spectrofluorometer FP-8300). The surface charge of the CNPs was determined using DLS/ZETA Sizer (Nano ZS, Malvern® Instruments).

### Statistical analysis

One-way Analysis of Variance (ANOVA) was carried out to determine the significant differences between the control and CNPs-supplemented treatments for the character’s primary root length and leaf area, followed by Duncan’s multiple range test using IBM SPSS Statistic version 22.

## Results

We have synthesized the carbon nanoparticles from *B. spectabilis, B. alba*, and *B. buttiana*, characterized the particle, and studied the effect of these particles on the vegetative growth of the *A. thaliana* as illustrated in **Fig.1**.

**Fig. 1.**
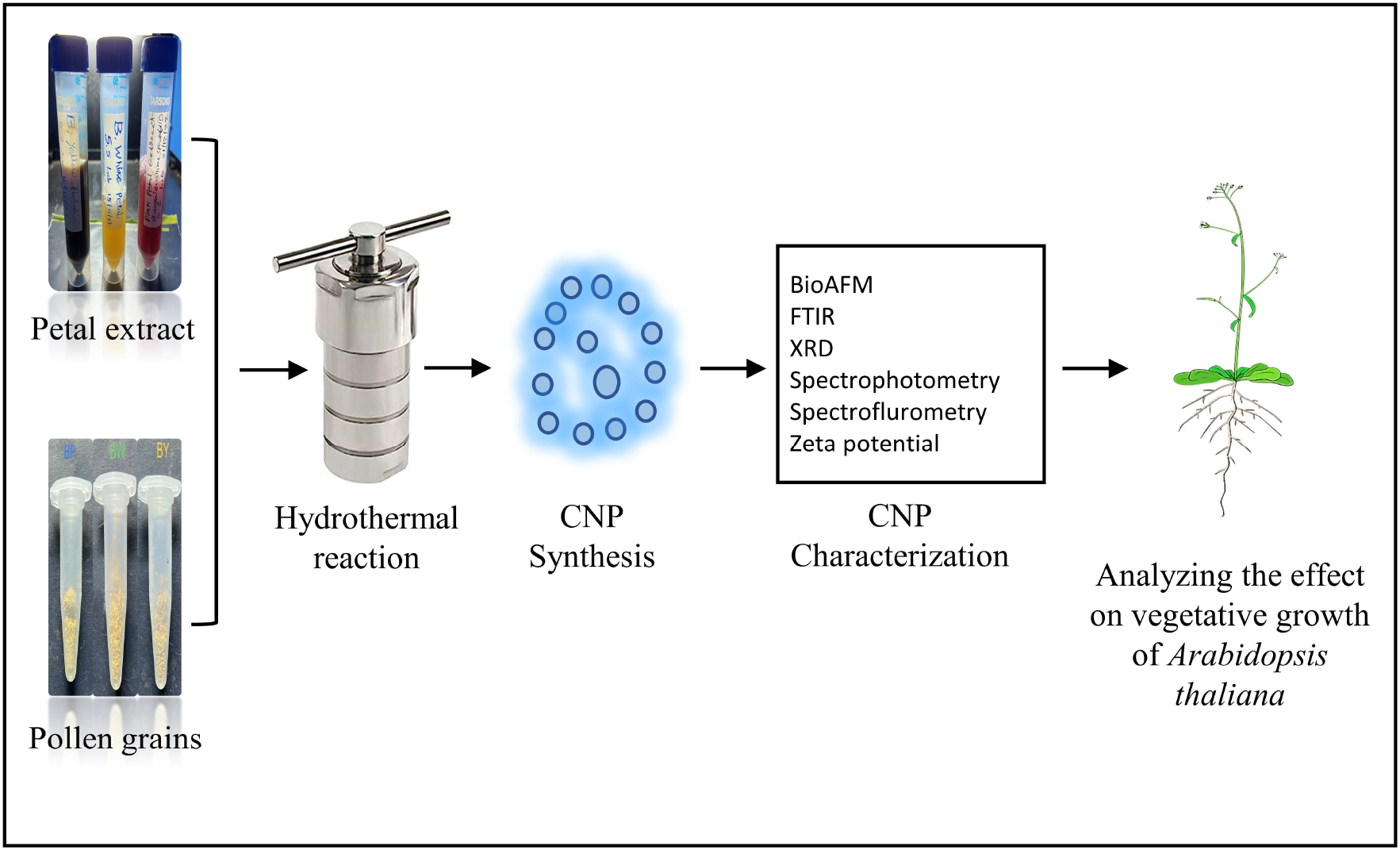
Synthesis of bract extract and pollen grains-derived carbon nanoparticles (CNPs) from *Bougainvillea* species and analyzing their effect on the vegetative growth of *A. thaliana*. CNPs were synthesized from the three *Bougainvillea* species, namely, *B. spectabilis, B. alba*, and *B. buttiana* using the hydrothermal method. CNPs were characterized and tested for their effect on the vegetative growth of *A. thaliana* plants.

### Pollen grains of *Bougainvillea* species have distinct morphological characteristics

The species *B. spectabilis, B. alba*, and *B. buttiana* have three bracts each and are red, yellow, and white, respectively **(Fig. 2A, B, and C)**. Pollen grains of the *Bougainvillea* species exhibited different morphological characteristics when analyzed under FE-SEM. *B. spectabilis* have a sub-oblate spheroidal shape **(Fig. 2D)**, *B. alba* has a spheroidal to prolate shape **(Fig. 2E)**, and *B. buttiana* a unique spheroidal shape (**(Fig.2F)** . *Bougainvillea* species also displayed different pollen exine coat patterns, where the exine of *B. spectabilis* **(Fig. 2G)** had a reticulum shape and round germ pore. The *B. alba*, on the other hand, had a thinner exine **(Fig. 2H)** compared to the other two species. However, *B. buttiana* exhibited an approximately similar pattern to *B. alba* without the germ pore but had thicker exine than *B. alba* (**Fig. 2I)**.

**Fig. 2.**
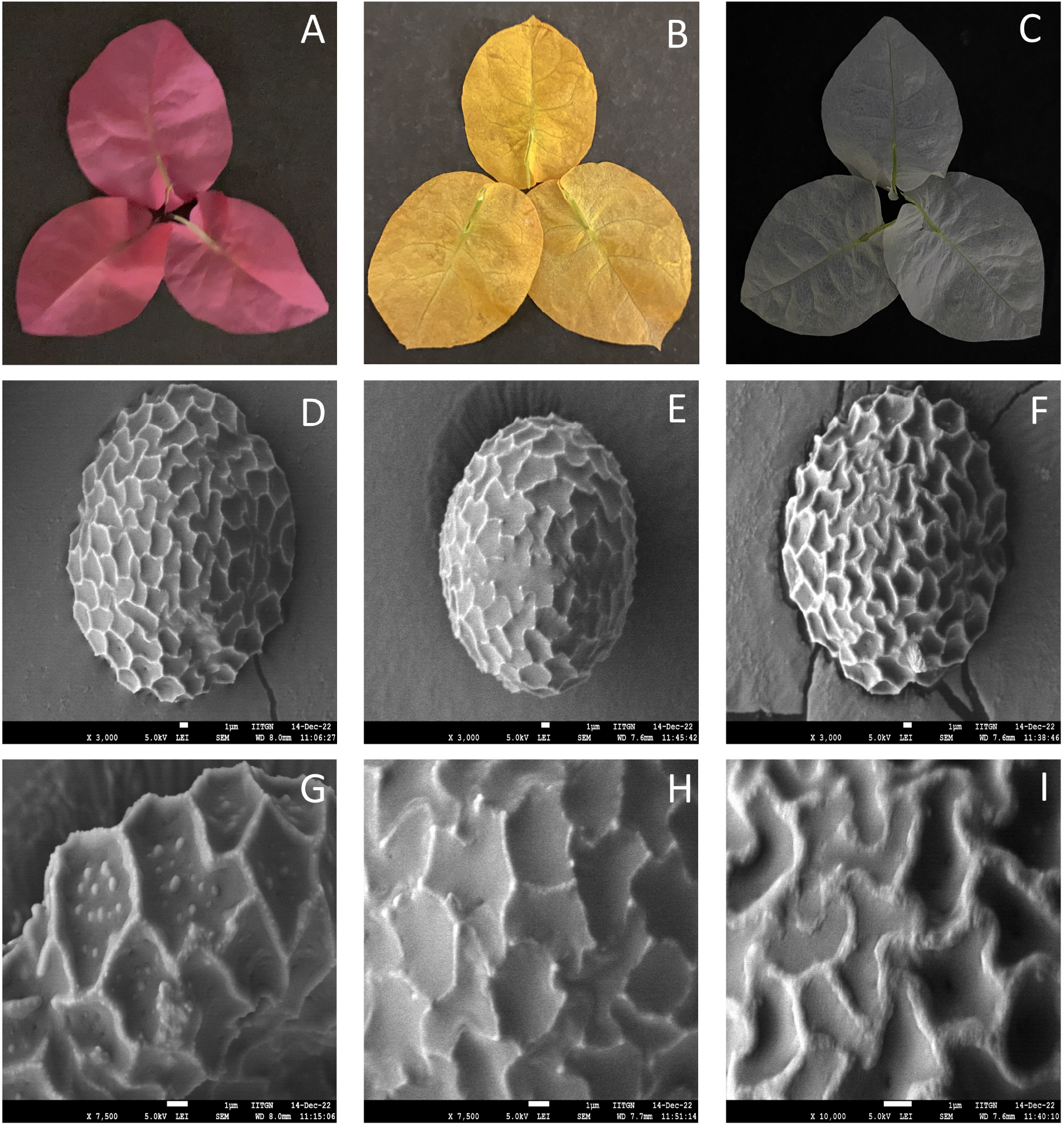
Morphological characteristics of *Bougainvillea* species. Images for (A) *B. spectabilis* bracts, (B) *B. alba* bracts, and (C) *B. buttiana* bracts were captured using a Camera. Images for (D) *B spectabilis* pollen grains, (E) *B. alba* pollen grains, (F) *B. buttiana* pollen grains, (G) *B. spectabilis* pollen exine coat, (H) *B. alba* pollen exine coat, and (I) *B. buttiana* pollen exine coat of were captured using FE-SEM. (D) to (F) were captured at 3000X, while images (G) and (H) were captured at 7500X, and (I) was captured at 10,000X. FE-SEM images were captured at LEI of 5.0 kV. The scale bar for the FE-SEM images corresponds to 1 µm.

### CNPs synthesized from *Bougainvillea* bracts and pollens have a size of less than 100 nm

The CNPs analyzed by Bio-AFM imaging revealed that the CNPs derived from bract extract of three different species of *Bougainvillea* had a size distribution of 83.49 nm ± 3.34 nm, 49.32 nm ± 1.48 nm, and 36.13 nm ± 1.66 nm for Bs-PET, Ba-PET, and Bb-PET, respectively **(Fig. 3A, B, and C, and Supplementary Fig. S2A, B, and C)**. The multipurpose X-ray diffraction (M-XRD) profile of bract extract CNPs, namely Bs-PET, Ba-PET, and Bb-PET, exhibited a broad peak centered around 23.7° (2θ), 18° (2θ), and 22° (2θ), respectively (**Fig. 2D, E, and F**). The corresponding *d*-spacing values for Bs-PET, Ba-PET, and Bb-PET are 3.75 Å, 4.92 Å, and 4.04 Å, respectively **(Fig 3D)**. The FTIR analysis of the bract extract CNPs revealed the presence of different functional groups on the CNP’s surface. The broad absorption band at 2960 cm^-1^ and 3025 cm^-1^ for Bs-PET and Ba-PET, respectively, indicated the stretching vibrations of the O-H group **(Fig. 3E and F)**. This was identified as a chemical with a hydroxyl group (-OH), which suggests that the surface still contains a significant amount of residual hydroxyl groups. The stretching vibrations of the O-H and N-H groups, which were described as compounds containing amine salt and a carboxylic acid, respectively, are represented by the absorption band centered at 2977 cm^-1^ for Bb-PET **(Fig. 3G)**. Furthermore, the three-bract extract CNPs Bs-PET, Ba-PET, and Bb-PET exhibited the bending vibration bands at 1489 and 1334 cm^-1^, 1487 and 1333 cm^-1^, and 1488 and 1333 cm^- 1^, respectively, representing the presence of C-H and O-H bands. In addition, the typical peaks at 1566 cm^-1^and 1116 cm^-1^of Bs-PET, 1567 cm^-1^and 1116 cm^-1^ of Ba-PET, and 1567 cm^-1^ and 1115 cm^-1^ of Bb-PET observed can be attributed to the stretching vibrations of C=C and C-N, respectively, representing the cyclic alkene and amine classes. These results suggest that the synthesized CNPs were N and S-doped (Sharma et al. 2019). The broad peak values in the X-ray diffraction profile indicated that the CNPs derived from the bracts of *Bougainvillea* were amorphous in nature. Nevertheless, the Zeta potential analysis of bract extract-derived CNPs had a negative charge, with values of -9.61 mV, -13.0 mV, and -6.47 mV for Bs-PET, Ba-PET, and Bb-PET, respectively.

**Fig. 3.**
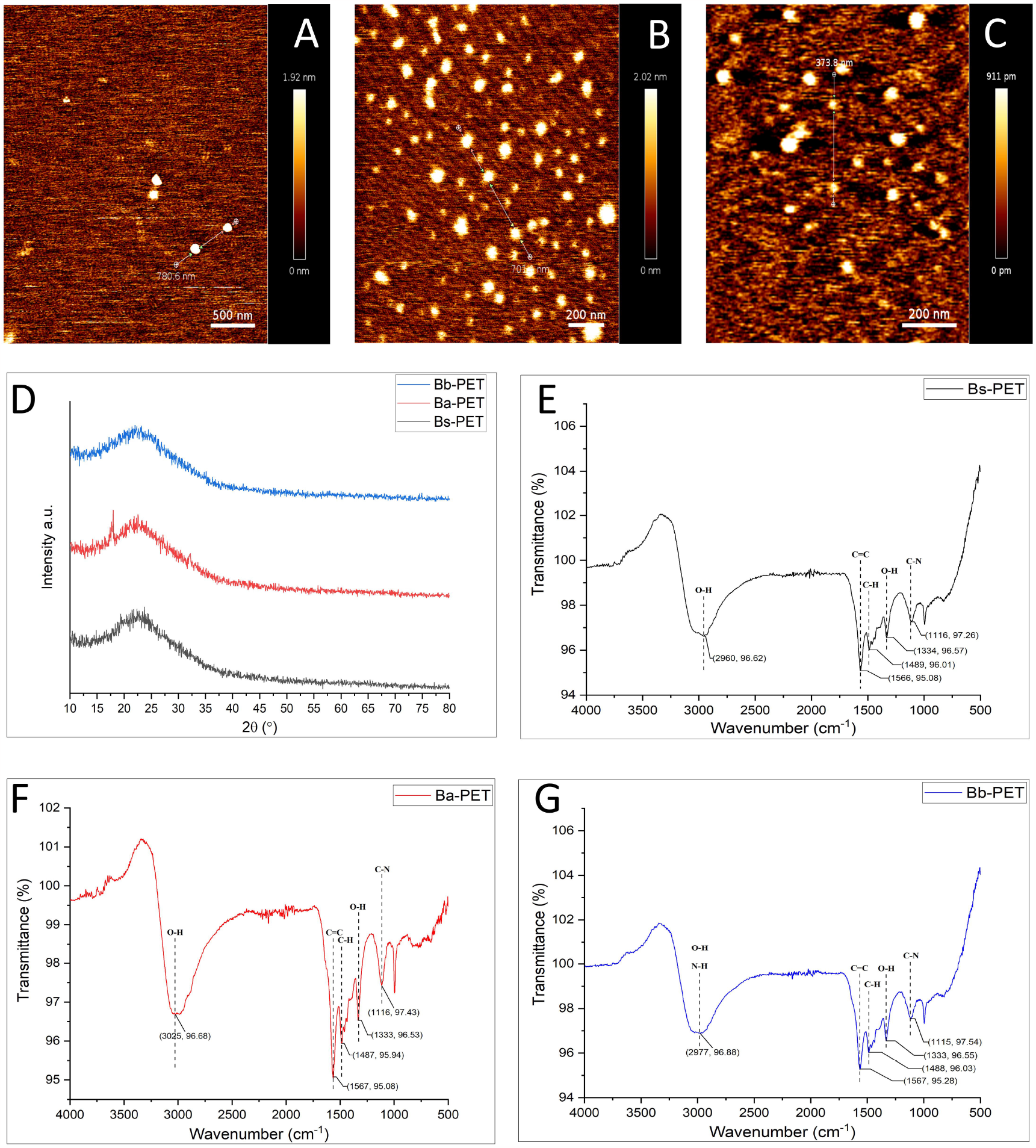
Characterization of carbon nanoparticles synthesized from bract extract of *Bougainvillea* species. The size of the bract extract-derived CNPs (A) Bs-PET, (B) Ba-PET, and (C) Bb-PET synthesized from *B. spectabilis, B. alba*, and *B. buttiana* was analyzed using Bruker® Nano wizard Sense AFM. The scale bar of Bio-AFM images (A) and (B) were 500 nm, and (C) was 200 nm. (D) M-XRD analysis of Bs-PET, Ba-PET, and Bb-PET was performed using Rigaku® SmartLab 9KW. M-XRD profile of CNPs was recorded with the scan rate of 4^°^ per minute in the range of 10^°^ to 80^°^ of 28 X-ray diffraction angle. (E), (F) and (G) represent the FTIR analysis of Bs-PET, Ba-PET, and Bb-PET, respectively, using Perkin Elmer® Spectrum Two FTIR Spectrometer. Graphs for FTIR and MXRD were plotted by using Origin 2017 version.

On the other hand, pollen-derived CNPs exhibited sizes of 59.96 nm ± 3.92 nm, 53.74 nm ± 2.32 nm, and 23.23 nm ± 0.95 nm for Bs-POL, Ba-POL, and Bb-POL, respectively **(Fig. 4A, B and C, and Supplementary Fig. S2D, E and F)**. Similarly, the M-XRD profile of pollen-derived CNPs Bs-POL, Ba-POL, and Bb-POL exhibited a broad peak centered around 23.3° (2θ), 24.25° (2θ), and 22.45° (2θ), respectively **(Fig. 4D)**. The corresponding d-spacing of Bs-POL, Ba-POL, and Bb-POL was found to be 3.81 Å, 3.67 Å, and 3.96 Å, respectively. FTIR profile of pollen-derived CNPs revealed that the typical peaks of Bs-POL and Ba-POL are respectively located at 2987 cm^-1^, 1572 cm^-1^, 1045 cm^-1^, and 2990 cm^-1^, 1575 cm^-1^, and 1045 cm^-1^, respectively **(Fig. 4E and F)**. These peaks correspond to the stretching vibrations of the C-H group (alkane), C=C group (cyclic alkene), and C-N group (amine classes). In contrast, the peak at 1416 cm^-1^ of Ba-POL represents the bending vibrations of the C-H group (alkane)and O-H group (carboxylic acid and alcohol classes). Moreover, the FTIR profile of Bb-POL represents that the typical peaks at 3335 cm^-1^ correspond to the stretching vibrations of the N-H (primary aliphatic amine and secondary amine) group **(Fig. 4G)**. The peaks at 2882 cm^-1^, 1973 cm^-1^, and 1054 cm^-1^, can be attributed to the stretching vibrations of the C-H (alkane), C=C=C (alkene), and C-N (amine) groups, respectively. The FTIR profile pollen-derived CNPs exhibited variation in the functional group and other groups on the surface of the CNPs. It may be because of the changes in morphology and other pollen content. The Zeta potential profile CNPs were -15.7 mV, -7.01 mV, and -10.2 mV for Bs-POL, Ba-POL, and Bb-POL, respectively. This negative charge represents good stability and low electrostatic repulsion between CNPs. The recorded negative value also indicates the ability to safely interact with cells during optical imaging, allowing for biomedical use (Mohiuddin et al. 2022).

**Fig. 4.**
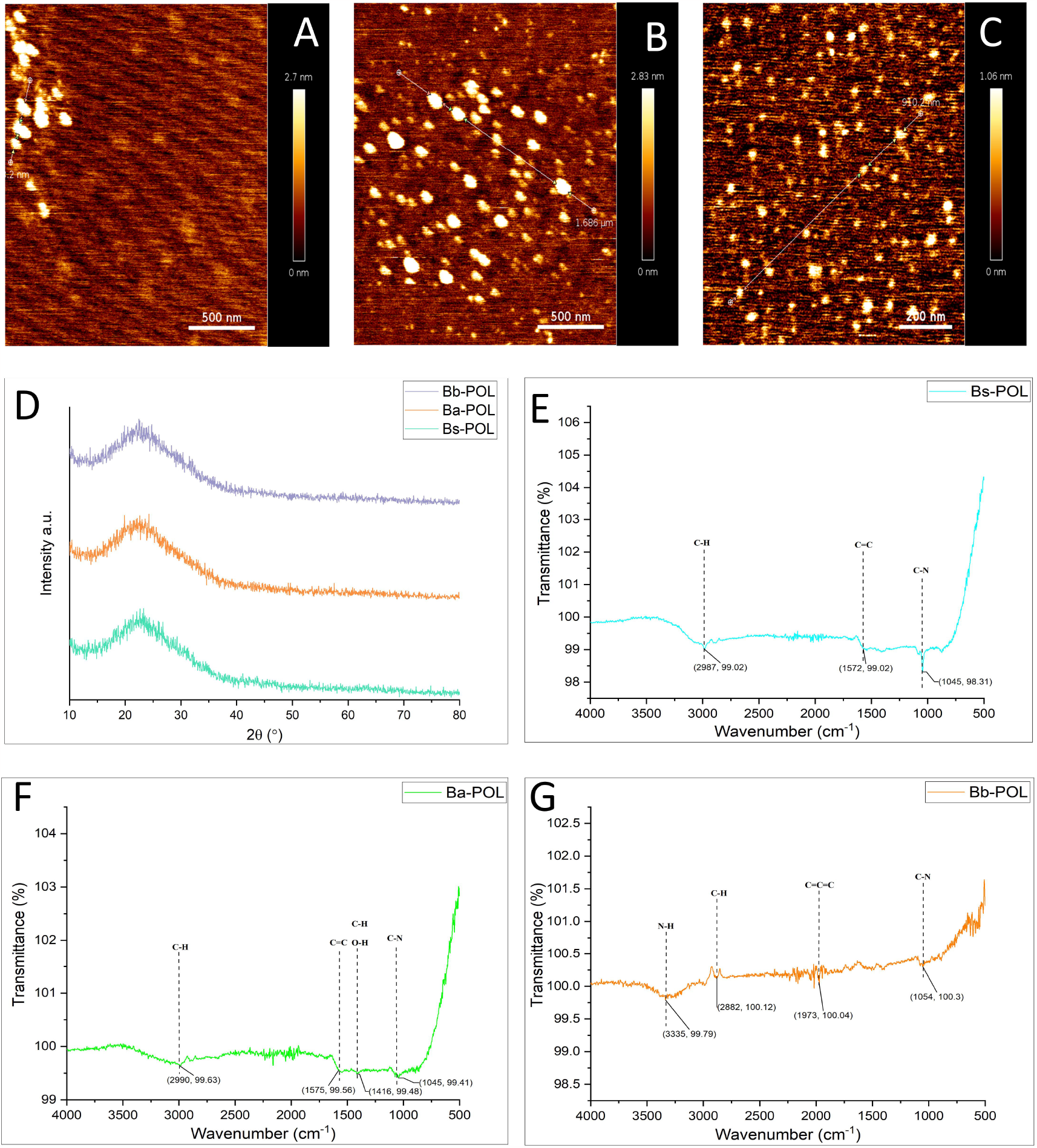
Characterization of synthesized carbon nanoparticles from the pollen of *Bougainvillea* species. The size of the pollen-derived CNPs (A) Bs-POL, (B) Ba-POL, and (C) Bb-POL synthesized from *B. spectabilis, B. alba*, and *B. buttiana*, respectively, were analyzed using Bruker® Nano wizard Sense AFM, with the scale bar 500 nm for (A) and (B) and 200 nm for (C). (D) M-XRD analysis for Bs-POL, Ba-POL, and Bb-POL, respectively, was carried out using Rigaku® SmartLab 9KW, with a scan rate of 4^°^ per minute and the range of 10^°^ to 80 ^°^ of 28 X-ray diffraction angle. (E) Bs-POL, (F) Ba-POL, and (G) Bb-POL represents the functional group analysis using Perkin Elmer® Spectrum Two FTIR Spectrometer. Graphs were plotted by using Origin 2017 version.

### CNPs emitted blue to purple-blue fluorescence under UV

The UV-visible absorption spectrum of bract extract CNPs displayed a noticeable peak at a wavelength between 310 and 320 nm in the lower-energy region between 300 and 600 nm **(Fig. 5A)**. This peak corresponds to the surface states with the presence of C=N, C=O, and C-O structures, as well as the n-π* transitions of the C=O bonds. The photoluminescence (PL) spectra of synthesized bract extract CNPs demonstrated a wide band emission of 250 nm to 500 nm for the three-bract extract CNPs. Under natural light or daylight conditions, the synthesized bract extracts CNPs Bs-PET, Ba-PET, and Bb-PET displayed a dark yellow colour in an aqueous solution. These CNPs emit purplish blue colour fluorescence under UV excitation **(Figure 5A)**. The highest fluorescence intensity was observed for Bs-PET, Ba-PET, and Bb-PET at 450 nm, 427 nm, and 394 nm, respectively when photoexcited at 370 nm, 360 nm, and 330 nm for bract extract-derived CNPs **(Figure 5B, C, and D)**.

**Fig. 5.**
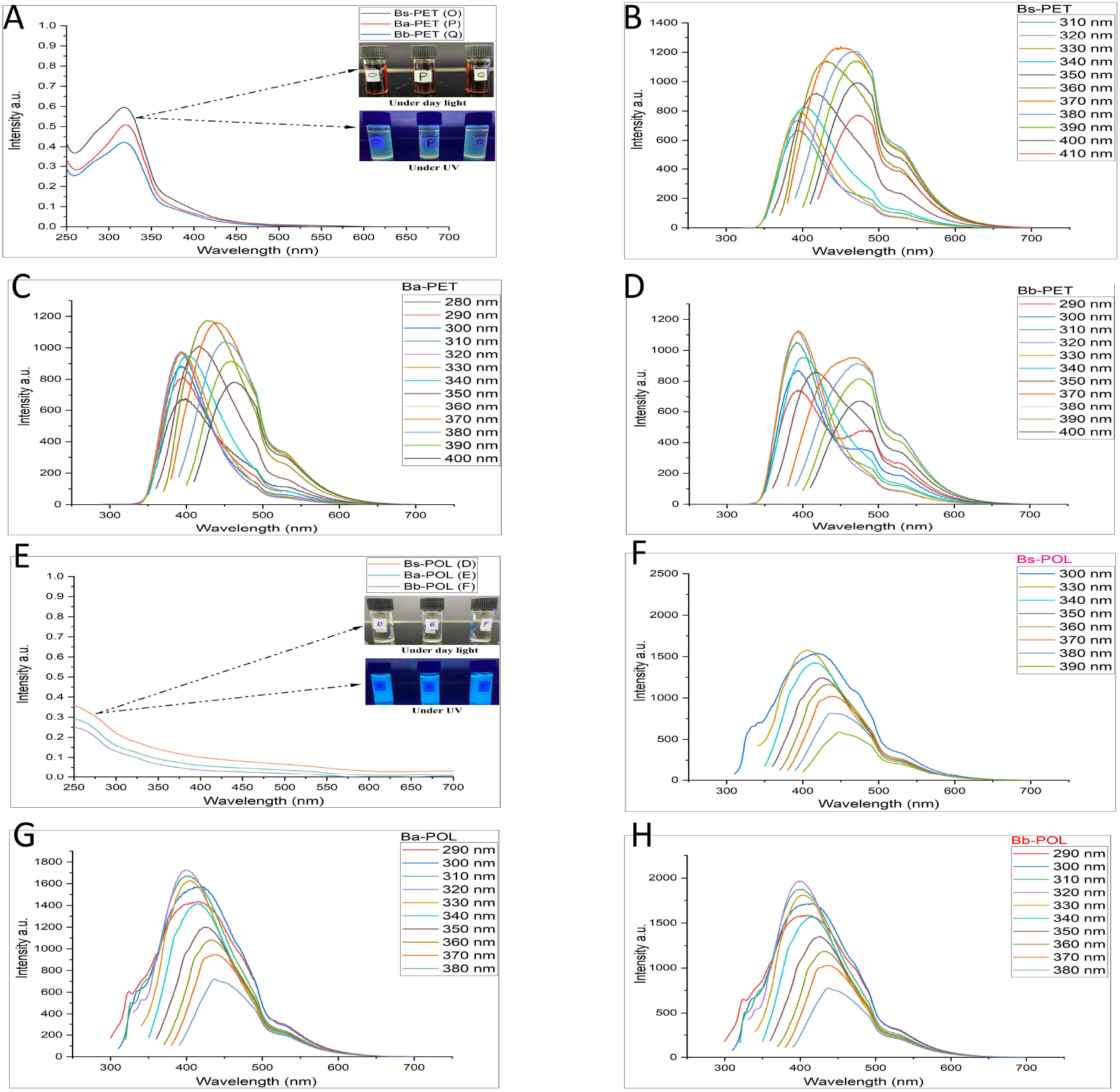
Characterization of optical properties of carbon nanoparticles synthesized from *Bougainvillea* species. Spectrophotometric analysis of bract extract-derived CNPs (A) Bs-PET, (B) Ba-PET, and (C) Bb-PET. The synthesized CNPs from Bs-PET, Ba-PET, and Bb-PET are labelled as samples 0, P, and Q, respectively. Spectrofluorometric analysis on CNPs, (B) Bs-PET, (C) Ba-PET, and (D) Bb-PET from species *B. spectabilis, B. alba*, and *B. buttiana*, respectively, was performed using Analytika Jena SPECORD® 210 PLUS spectrophotometer and Jasco® spectrofluorometer FP-8300. Spectrophotometric and spectrofluorometric analyses were conducted on pollen-derived CNPs: (D) Bs-POL, (E) Ba-POL, and (F) Bb-POL, with the corresponding synthesized CNPs labeled as D, E, and F, respectively. The analysis also included spectrofluorometric analysis of (E) Bs-POL, (F) Ba-POL, and (G) Bb-POL in the range of 250 nm to 700 nm. Graphs were plotted by using Origin 2017 version.

In pollen-derived CNPs (Bs-POL, Ba-POL, and Bb-POL), the UV-visible absorption spectra displayed a prominent peak in the higher-energy UV range between 250 to 280 nm. Pollen-derived CNPs show broad emission from 350 nm to 480 nm, and under natural light or daylight conditions, pollen-derived CNPs are transparent in colour and emit strong blue fluorescence under UV excitation **(Fig. 5E)**. The fluorescence intensities of Bs-POL, Ba-POL, Bb-POL peak at 406 nm, 399 nm, and 400 nm, respectively, when photoexcited at 330 nm, 320 nm and 320 nm **(Fig. 5F, G and H)**.

### CNPs increased the leaf area but suppressed the primary root growth

The synthesized CNPs tested on *A. thaliana* displayed a stimulatory effect on the leaf area of plants in a dose-dependent manner after seven days of the treatments with CNPs (**Fig. 6 and Supplementary Table S1A to C**). The leaf area increased to the extent of 43.8% at a concentration of 5 mg/ml in Ba-POL-supplemented treatments (**Fig. 6C)**. A similar effect on the significant increase in leaf area at the concentrations of 2.5 mg/ml and 10 mg/ml was also observed in Ba-POL-supplemented treatments (**Supplementary Table S1A to C**). These results suggest that the CNPs synthesized from *B. alba* have a significant effect on the increase in leaf area which can be observed at different concentrations.

**Fig. 6.**
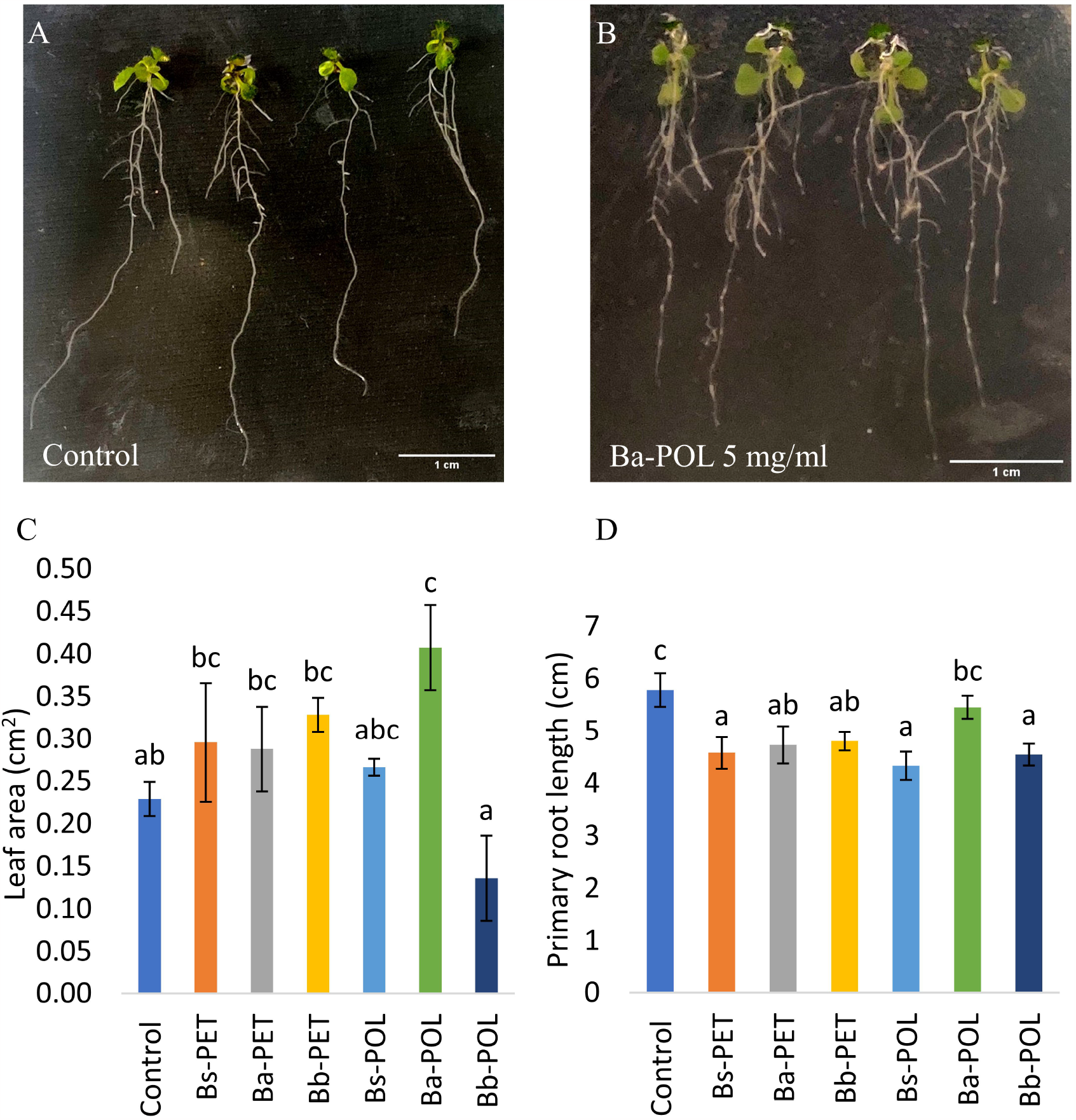
Plant growth assay of carbon nanoparticles (CNPs) derived from the bract extract and pollens grains of *Bougainvillea* species on *Arabidopsis thaliana*. (A) and (B) represent the control and 5 mg/ml *B. alba* pollen-derived CNPs (Ba-POL) treatment. The leaf area (cm^2^) and primary root lengths were measured using Image J after seven days of the treatments. (C) and (D) represent the increase in the leaf area and suppression in the primary root length in response to a 5 mg/ml concentration of bract extract-derived CNPs (Bs-PET, Ba-PET B, and Bb-PET), and pollen-derived CNPs (Bs-POL, Ba-POL, and Bb-POL) in the species *B. spectabilis, B. alba* and *B. buttiana*, respectively. Control represents normal growth without supplementation of CNPs. Significant differences between the treatments were analyzed using one-way ANOVA at p≤S0.05, followed by Duncan’s multiple-range test. Statistical analysis was carried out using IBM SPSS Statistic version 22.

After day seven of the treatment, CNPs tested for the effect on the primary root growth demonstrated significant inhibition in the primary root growth, except for Ba-POL-synthesized CNPs. The highest suppression of 26% in the root growth was found in the treatments supplemented with Bs-POL, followed by 20.8% suppression in Bs-PET. Though a significant reduction in other treatments for the primary root growth was observed, the Ba-POL-supplemented treatment suppressed the root growth by only 5.6% (**Fig. 6A, B, and D**). These results suggest that CNPs synthesized from *B. alba* had a minimal effect on the suppression of PR growth at a concentration of 5mg/ml, whereas CNPs synthesized from *B. spectabilis* had exactly the opposite effect and suppressed the root growth at similar concentrations (**Supplementary Table S1D to F**).

## Discussion

### Bract and pollen-derived CNPs have a unique composition

*Bougainvillea* species are known to exhibit various characteristic properties and have been successfully utilized in horticulture and medicinal applications. Different chemical compounds have been identified in the flower and bract of these species, including fatty acids, alcohols, phenols, volatile compounds, terpenes, and other compounds (Abarca-Vargas and Petricevich 2018). We have reported the synthesis of the nanoparticles from the bract extract and pollen grains of the three *Bougainvillea* species (*B. spectabilis, B. alba*, and *B. buttiana*), namely Bs-PET, Ba-PET, Bb-PET, Bs-POL, Ba-POL, and Bb-POL. Similar studies have been conducted on the synthesis of carbon quantum dots (CQDs) from rose bracts (Sharma et al. 2019). Nevertheless, the bract-derived nanoparticle sizes are larger than the rose quantum dots and were mainly composed of carbon. Both these quantum dots emit blue coloration under ultraviolet illumination, are negatively charged, and have amorphous nature. Zhang et al. 2015, also reported that the synthesized fragrant nitrogen-doped CQDs from bee pollen are also amorphous. Proteomic studies on the different stages of anther and pollen have been conducted in the plant species like *Arabidopsis* and rice (Imin et al. 2001; Holmes-Davis et al. 2005; Noir et al. 2005). In *Arabidopsis* mature pollen, the proteomic analysis identified 135 unique proteins involved in various processes such as metabolism, cell structure, and protein synthesis (Holmes-Davis et al. 2005; Noir et al. 2005). We have synthesized the nanoparticles from the pollen grains of three *Bougainvillea* species, which have different structures of exine and potentially distinct protein compositions. This feature of pollen can be utilized to investigate the effect of pollen-derived CNPs from different species. The size of the CNPs synthesized from different sources is in the range of 10-100 nm. CNPs synthesized from nickel and carbon-encapsulated nickel had the size of nanoparticles in the range of 10 to 50 nm, whereas for gold nanoparticles the sizes were in the range of 14, 30, 50, 74, and 100 nm (Cheng et al. 2006; Chithrani et al. 2006). The synthesized CNPs were also in the same ranges and had a size of less than 100 nm.

### Source and method of synthesis affect CNP characteristics

Studies have documented the synthesis of CNPs using date palm fronds as a source. For instance, Kavitha and Kumar (2018) used hand grinding to carbonize the fronds of date palms, resulting in CNPs with an average size of 32 nm. The lignin nanoparticles were synthesized using date palm tree biomass by hydrothermal method, yielding CNPs with diameters ranging from 5 to 15 nm (Athinarayanan et al. 2022). XRD analysis revealed that these particles were crystalline. This suggests that the properties of CNPs depend on the source of the biomass and the synthesis method employed. The presence of hydrophilic groups present in CNPs could provide insight into the mechanics of luminescence and contribute to the development of more environmentally friendly products for use in biological settings (Liu et al. 2014). The recorded negative value for the Zeta potential also indicates the ability of CNPs to interact safely with cells during optical imaging, making them suitable for biomedical use (Mohiuddin et al. 2022). CNPs synthesized in this study were amorphous in nature and had a negative Zeta potential value, which provides an opportunity to test these nanoparticles for different applications.

### CNPs concentration is the critical parameter to test the effect on plants

The effect of the nanoparticles on plant growth and development and stress responses have been widely studied (Kumar et al. 2013; Dimkpa et al. 2019; Ferrari et al. 2021). The positive effect of the nanoparticles on plant growth and development depends on various factors such as concentration, source, size, and time of nanoparticle application to the crop (Rubilar et al. 2013). The gold-derived nanoparticles AuNP have been shown to promote root growth and decrease the stress response in *Arabidopsis* seedlings, with 10mg/ml as the most significantly affecting concentration (Ferrari et al. 2021). In another study, the gold-derived nanoparticles with 24 nm size at 10 µg/ml concentration significantly improved the yield, germination rate, vegetative growth, and free radical scavenging in *Arabidopsis* (Kumar et al. 2013). These studies provide evidence that concentration is the critical factor in studying the effect on the plant’s growth. This study has investigated the effects of the different concentrations of nanoparticles and demonstrated the concentration of 5mg/ml as the most effective in increasing leaf surface area in *Arabidopsis*. However, similar concentrations also had the inverse effect on the primary root growth. CNPs synthesized from the white bract *Bougainvillea*, Ba-POL increased the leaf area significantly at different concentrations, with an increase of 43.8% at 5 mg/ml concentration. Interestingly, the white-bract *B. alba* has a thinner exine compared to the other two species, and FTIR analysis showed the peak at 1416 cm^-1^ for this species, which indicates the presence of alkane, carboxylic acid, and alcohol groups. These results suggest that CNPs concentration plays a critical role in the phenotypes of the plants and the pollen from the same genus species have very unique compositions and can have different effects on phenotypes.

### CNPs effect on vegetative growth may be due to interference in the auxin signaling

Hormonal signalling has a crucial role in plant growth and development. The hormone auxin is synthesized in the shoot apex and transported to the root tips, regulating cell elongation and divisions in plants. We hypothesized different possibilities for the effect on the phenotypes observed upon treatment with CNPs. The first possibility is that the CNPs synthesized in this study may interfere with the auxin signalling pathway, as well as transport and distribution within the plant, which results in the accumulation of auxin in the shoot region to promote shoot growth and make it unavailable to roots and inhibit root growth. A similar effect on the CNPs on the root growth inhibition in the primary root to the extent of 55% was observed in *Arabidopsis* Col-0 wild-type seedlings, and it was found that carbon dots can enter the meristem, elongation, and root hair zone of the primary root. In a similar study, the use of the DR5:GUS line also suggested that the root meristem development suppression by CDs may be due to the decrease in the auxin concentration at the root tip. Nevertheless, the decrease in the gene expression levels of seven auxin biosynthesis genes was also observed when treated with CDs (Yan et al. 2021). This suggests that a similar effect could also be found with the application of CNPs in this study. Ba-POL synthesized CNPs had less effect on the suppression of primary root growth compared to other synthesized CNPs at the concentration of 5 mg/ml, and significantly increased the leaf area at three different concentrations of 2.5 mg/ml, 5 mg/ml, and 10 mg/ml. This suggests that the 5 mg/ml concentration of Ba-POL is beneficial either way since it increases leaf area and has minimal effect on root growth suppression.

Interestingly, we have also observed this trend in the treatments supplemented with Ba-POL, where greater inhibition in the primary root growth was correlated with increased leaf area. However, this effect was maximum at a concentration of 5 mg/ml. This trend indicates the allocation and utilization of the auxin in the shoot growth (**Supplementary Table S1A to F**). The hormonal signalling pathways are complex, and the CNPs may interfere with multiple pathways including cytokinin which stimulates leaf expansion, and the abscisic acid (ABA) pathway, which also suppresses root growth. Moreover, ABA-mediated root growth inhibition occurs due to a reduction in the auxin level in roots, and its inhibitory effect is attributed to Reactive Oxygen Species (ROS) production (He et al. 2012). Hence, it is plausible that CNPs suppress root growth by inducing oxidative stress and generating (ROS) in plants, which, in turn, may have a compensatory response in plants, leading to resource allocation in plants to enhance the above-ground phenotypes, such as leaf area. Nevertheless, the comprehensive analysis of the molecular mechanism underlying the effect of CNPs on hormonal signalling during plant growth could be an interesting area that needs to be investigated further.

## Conclusion

In this study, we successfully synthesized the six different CNPs from bract extract and pollen grains of three different species of *Bougainvillea* using a single-step hydrothermal reaction. The spectrophotometric and spectrofluorimetric analyses confirmed that the CNPs displayed violet and blue emission when subjected to UV illumination. FE-SEM analysis of the pollen exine revealed the distinct characteristics of the exine of three *Bougainvillea* species. CNPs were amorphous and negatively charged. CNPS tested for the effect on the vegetative growth of *A. thaliana* and exhibited a leaf area-promoting response, and suppressed the primary root growth. Further investigations could be conducted to explore additional applications of these CNPs and consider their potential use as a spray in agriculture to enhance vegetative growth in plants.

## Acknowledgment

We thank the Indian Institute of Technology Gandhinagar for the internship opportunity for RB and the post-doctoral fellowship to NG. This work was supported by a DBT Ramalingaswamy Re-entry fellowship grant and a start-up grant from the Indian Institute of Technology Gandhinagar to SS.

## Author Contributions

SS conceived and designed the research. RB synthesized and characterized the CNPs and NG assisted in the characterization of CNPs. RB, SO, and CK performed experiments on the effect of CNPs on the leaf area and root growth in Arabidopsis. SS supervised the experiments. NG and SS analyzed the data. RB wrote the manuscript, and NG and SS proofread and edited the manuscript. JS and DB provided technical suggestions for experiments and shared reagents. All authors read, discussed, and approved the manuscript.

## Additional Information

### Competing Interests

The authors declare no competing interests.

## Notes

https://www.supplementaryfiles.com

### Competing Interest Statement

The authors have declared no competing interest.

