## Supplementary figures and images for "Novel carbon nanoparticles derived from *Bougainvillea* modulate vegetative growth in *Arabidopsis*"

### Supplementary Figure S1.tif

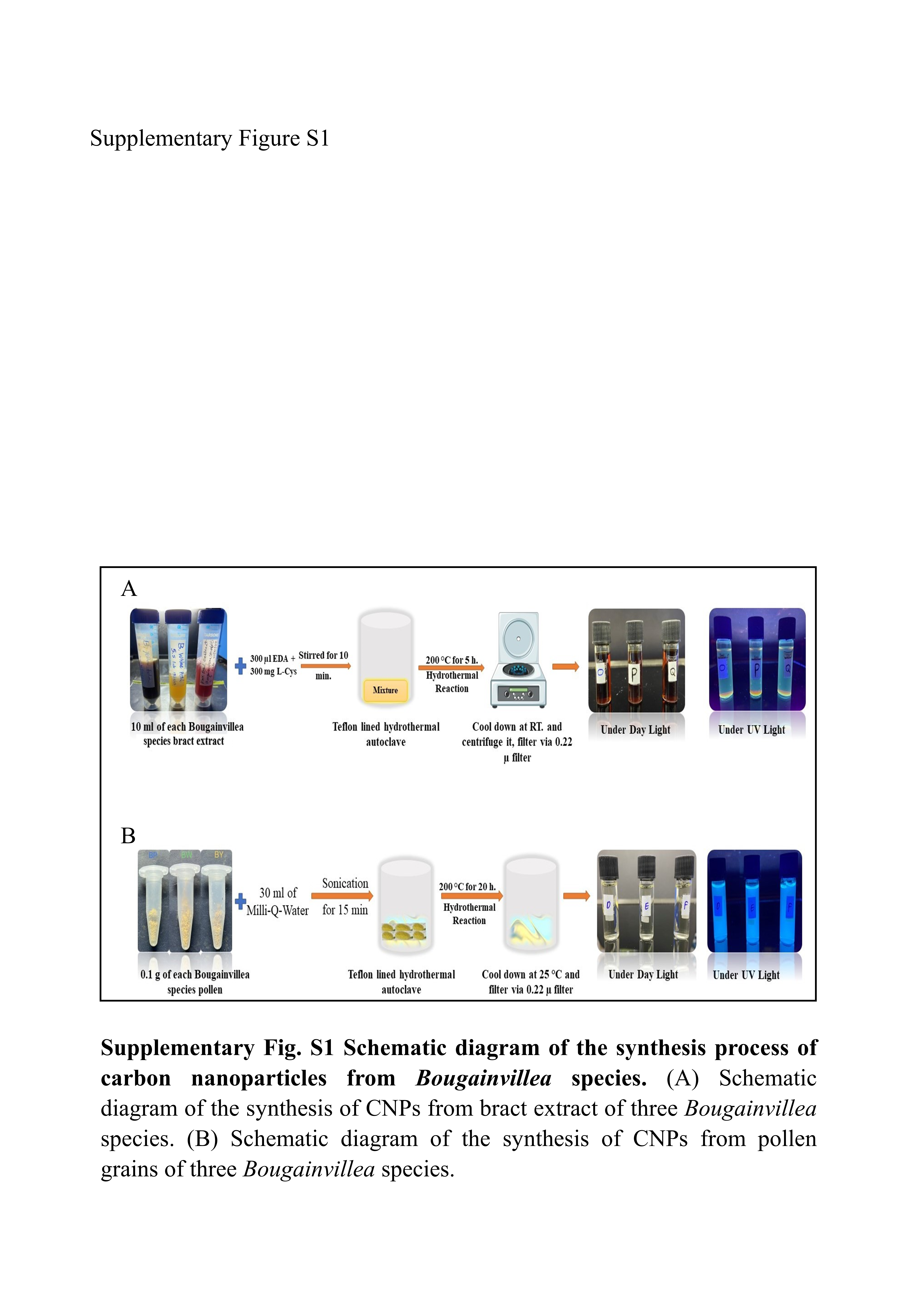

### Supplementary figure S2.tif

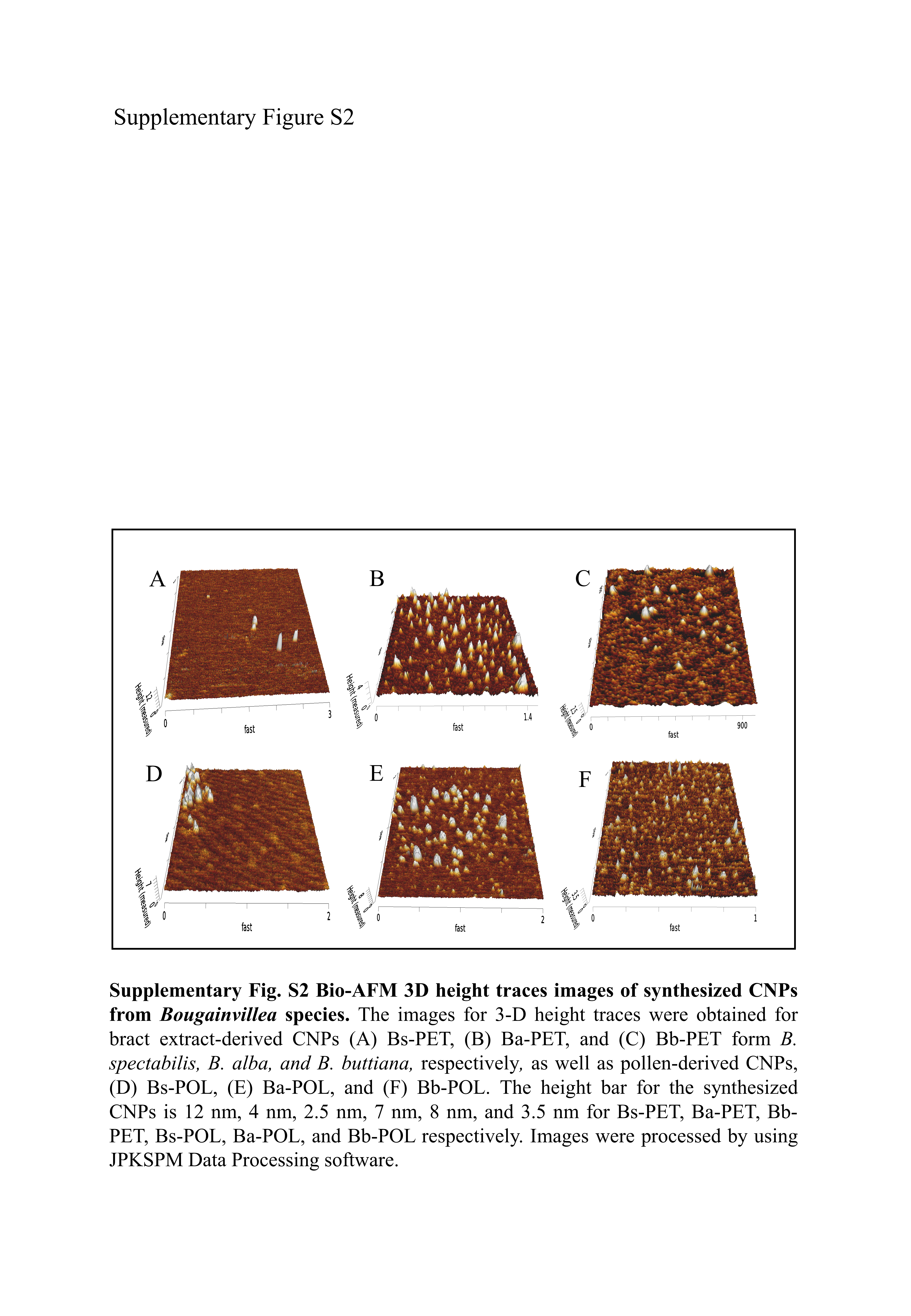
